# Sex and Alkyladenine DNA Glycosylase Expression are Key Susceptibility Factors for NDMA-induced Mutations, Toxicity, and Cancer

**DOI:** 10.1101/2025.05.13.653839

**Authors:** Jennifer E Kay, Joshua J Corrigan, Lindsay B Volk, Amanda L Armijo, Ilana S Nazari, Dorothea K Torous, Svetlana L Avlasevich, Robert G Croy, Dushan N Wadduwage, Stephen D Dertinger, John M Essigmann, Leona D Samson, Bevin P Engelward

## Abstract

*N*-Nitrosodimethylamine (NDMA) is present in food, water, and drugs and is considered a probable human carcinogen by the International Agency for Research on Cancer. The mechanism of action of NDMA involves the generation of carcinogenic methyl lesions such as 3-methyladenine (3MeA) on DNA bases. Alkyladenine DNA Glycosylase (AAG) removes 3MeA to initiate Base Excision Repair, leaving an intermediary lesion that is subsequently resolved by backbone cleavage, nucleotide insertion, and backbone ligation. The intermediate steps following lesion removal produce potentially toxic and mutagenic single-strand DNA breaks. Here, we explored differences between males and females regarding downstream DNA damage, toxicity, mutations and cancer arising from 3MeA in the livers of WT, *Aag*^-/-^, and *Aag*-overexpressing (*AagTg*) mice. We found that males were more susceptible to NDMA-induced mutations (WT and *Aag*^-/-^) and cancer (all genotypes). In contrast, *AagTg* females were more prone than males to micronucleus induction. As we showed in our prior analyses where data were pooled for males and females, *Aag*^-/-^ mice were significantly more susceptible to NDMA-induced mutations and cancer, and *AagTg* mice displayed significantly greater toxicity. Building on these findings, our analyses of sex-related differences show that *Aag* deficiency and maleness are both susceptibility factors for NDMA-induced liver cancer, while *Aag* overexpression drives toxicity, potentially with a greater effect on females. By assessing differences between males and females, this study reveals a deeper mechanistic understanding of the underpinnings for a well-known increased risk of liver cancer in men versus women by demonstrating a higher susceptibility of male mice to both mutations and cancer.

## INTRODUCTION

*N*-Nitrosodimethylamine (NDMA) is a potent carcinogen in animal models and is a Group 2A probable human carcinogen according to the International Agency for Research on Cancer (IARC)^1^. Humans are exposed to NDMA through widespread sources. NDMA can be found in drinking water as a consequence of commonly used water treatment processes^2–6^, as well as ineffectively contained hazardous waste^7–10^. NDMA can also be found in many foods, including processed meats, which are considered to be carcinogenic according to IARC^7–10^. Recently, NDMA has been found as a contaminant in numerous drugs that have since been recalled^11–15^. Consequently, NDMA is a significant public health concern. Here, we have explored differences in susceptibility to NDMA-induced DNA and tissue damage, mutations, and cancer for males versus females.

The main mode of action for NDMA is through the induction of mutagenic DNA lesions that ultimately drive cancer. Metabolism of NDMA via CYP2E1 leads to formation of the methyldiazonium ion, which is highly unstable and can react to add methyl groups on DNA bases, producing lesions such as 3-methyladenine (3MeA)^7^. In animal models, the risk of cancer for different tissues reflects their CYP2E1 metabolic capacity, and CYP2E1 is expressed at the highest levels in the liver^16^. We recently demonstrated the mutagenic potential of 3MeA *in vivo*^20^. To defend against 3MeA-driven mutations, 3MeA is removed by the Alkyladenine DNA Glycosylase (AAG; a.k.a., MPG)^21^, which initiates base excision repair (BER). Following excision of 3MeA by AAG, the resulting abasic site is processed by APE1 to create a nick in the DNA backbone 5’ to the lesion. In short-patch BER, DNA polymerase beta removes the abasic ribose and inserts a nucleotide. Finally, the nick is sealed by DNA ligase III. The BER process is highly coordinated to minimize the presence of toxic BER intermediates, such as requisite single-strand breaks^22^.

Unrepaired 3MeA can block replicative polymerases, leading to replication fork stalling and toxicity^23–25^. However, *in vitro* studies have demonstrated that some translesion synthesis (TLS) polymerases can bypass 3MeA ^25–28^, and our prior work showed that 3MeA is a point mutagen *in vivo*^20^. An alternative pathway for coping with replication-blocking lesions is to invoke homologous recombination (HR). For example, if the leading strand is stalled, DNA replication can continue by using the newly synthesized lagging strand as a template^23^. While HR is mostly error-free, misalignments can lead to insertions, deletions, and translocations. Consistent with the ability of replication blockage by 3MeA to cause point mutations and larger scale sequence rearrangements, AAG-deficient mice that are unable to repair 3MeA are highly susceptible to NDMA-induced liver cancer^20^. Conversely, mice overexpressing AAG have reduced levels of NDMA-induced cancer compared to wild-type (WT) mice^20^. However, *Aag*-overexpressing mice also suffer increased toxicity, presumably due to single strand breaks that are formed during BER. Thus, the fate of the cell following alkylation exposure depends on a balance between efficient AAG excision and coordinated downstream BER machinery to resolve DNA strand breaks.

Interestingly, the mutagenic and carcinogenic consequences of alkylating exposures depend not only on the levels of DNA repair enzymes, but also on sex. Generally, the incidence and mortality rate of non-reproductive cancers are elevated in males compared to females^29,30^. Indeed, the occurrence of hepatocellular carcinoma (HCC) is 2-3 times higher in males versus females^30^. Though lifestyle plays an intricate role in the elevated rates of cancer in men, intrinsic factors such as sex hormones greatly influence carcinogenesis, especially in a sexually dimorphic organ such as the liver. Sex also influences the tumorgenicity of environmental carcinogens^31^. In the context of nitrosamine exposure, several studies have demonstrated that females develop fewer liver tumors than males^18,32–34^.

We previously interrogated the biological consequences of NDMA exposure in mice lacking AAG expression (*Aag*^-/-^) or overexpressing AAG (*Aag* transgenic; *AagTg*), which are prone to accumulating either 3MeA lesions or BER intermediates, respectively. We analyzed a suite of endpoints spanning the days immediately after exposure to months later. By integrating analyses of DNA and tissue damage, animal lethality, mutations, and cancer, we were able to track the progression from exposure to disease. We found that unrepaired 3MeA (as in *Aag^-/-^*mice) is somewhat toxic but highly mutagenic, whereas excess strand breaks (as in *AagTg* mice) are poorly mutagenic and highly toxic^20^. In that study, we pooled results from males and females to specifically analyze the biological impacts of AAG expression. Here, we have taken that research a step further by revealing sex-specific differences between males and females. Males were generally more susceptible to NDMA-induced long-term liver damage and cancer in all genotypes, which is consistent with prior work^18,32–34^. Concordant with their increased susceptibility to cancer, males were also more vulnerable to sequence rearrangement mutations. These effects were primarily elevated in *Aag^-/-^* mice and suppressed in *AagTg* mice compared to WT. These studies reinforce a model wherein physiological conditions favoring mutations and disfavoring toxicity are highly carcinogenic, while physiological conditions that are less favorable to mutations and more favorable to toxicity are associated with suppression of cancer.

## MATERIALS AND METHODS

The data presented here are derived from male and female combined results presented in our previous publication Kay et al., 2021^20^, which contains details regarding materials and methods. Here, we briefly summarize experimental approaches and point the reader to our previous manuscript for details.

### Animals

The *RaDR^R/R^*;*gpt^g/g^, RaDR^R/R^*;*gpt^g/g^*;*Aag^−/−^,* and *RaDR^R/R^*;*gpt^g/g^*;*AagTg* mice utilized in this study were on a C57BL6 background^20^. Standard chow and drinking water were provided *ad libitum* in an AAALAC-certified animal care facility. All animal procedures were performed in accordance with the NIH Guide for the Care and Use of Laboratory Animals and approved by the Massachusetts Institute of Technology Committee on Animal Care.

### NDMA Treatment

Mice were administered two intraperitoneal injections of NDMA diluted in saline, 3.5 mg/kg NDMA at 8 days of age and 7 mg/kg NDMA at 15 days of age for a total dose of 10.5 mg/kg NDMA as previously described^20^. Control mice were treated with equal volumes of saline at the respective timepoints. Depending on the assay, mice were euthanized and necropsied at either 24 hours, 48 hours, 10 weeks, or approximately 10 months post-second injection. At the time of necropsy, all visible macroscopic surface lesions on the liver were enumerated. Timepoint details are included in the figure legends.

### Sequence Rearrangement Mutation Analysis (RaDR)

Sequence rearrangement mutations were analyzed using the Rosa26 Direct Repeat (RaDR) transgenic mouse model^20^. Briefly, the left liver lobe was freshly excised from mice and put into ice-cold PBS with 0.01% trypsin inhibitor. The lobe was then placed on a slide and the dorsal surface was imaged with a 2x objective in the FITC channel. A machine learning algorithm was utilized to quantify the number of fluorescent foci within the lobe, and foci counts were normalized to the area of the lobe^20^.

### Point Mutation Analysis (Gpt**Δ** assay)

Liver samples were flash frozen in liquid nitrogen and stored at −80°C. λ-EG10 phage containing genomic DNA was extracted from the liver samples, then packaged and infected into *Escherichia coli* YG6020^35^. These bacteria were cultured on chloramphenicol and 6-thioguanine selective media plates. Mutations in the *gpt*Δ gene were measured by the number of 6-thioguanine resistant colonies.

### **γ**H2AX Staining and Analysis

Liver samples were formalin fixed, paraffin embedded, and sectioned at 4-μm thickness prior to staining. Liver sections were deparaffinized, rehydrated, and subjected to heat-induced epitope retrieval. Following blocking, the sections were incubated with γH2AX antibody (1:200; Cell Signaling Technologies) overnight at 4°C. Washed sections were incubated with AlexaFluor 488 secondary antibody (1:400; Invitrogen) then mounted with ProLong Gold AntiFade Mountant with DAPI (ThermoFisher). Stained sections were imaged with a 60x objective under DAPI and FITC filters. Percentages of nuclei with 5 or more γH2AX foci, pan-nuclear γH2AX staining, and super-bright pan-nuclear staining were analyzed blinded with ImageJ^20^.

### Micronucleus (MN) Assay

Livers were excised and immediately placed in ice-cold Liver Preservation Buffer, then shipped to Litron Laboratories. Livers were processed for flow cytometry as described in Avlasevich et al., 2018^36^, and analyzed for SYTOX nuclear stain (488 nM excitation) and anti-Ki67-eFluo^R^ 660 (633 nM excitation) on a FACSCanto™ II flow cytometer (BD Biosciences, San Jose, CA)^20^. Percent of micronucleated hepatocytes was quantified as number of MN / number of nuclei x 100.

### Cleaved Caspase-3 Staining and Analysis

Liver sections were prepared as in γH2AX Staining. Following heat-induced epitope retrieval, slides were blocked with Peroxidazed 1 (Biocare Medical) then Background Sniper (Biocare Medical) prior to incubation with cleaved caspase-3 antibody (Cell Signaling Technology). Washed sections were incubated with Rabbit-on-Rodent HRP Polymer (Biocare Medical), then washed and incubated with DAB (Betazoid DAB Chromagen kit, Biocare Medical), then washed and counterstained with hematoxylin. Stained slides were scanned at 40x and analyzed for percentage of apoptotic events using QuPath software^20^.

### Histological Analysis

Formalin-fixed paraffin-embedded liver samples were sectioned and stained with hematoxylin and eosin (H&E). H&E liver sections were scored by a board-certified veterinary pathologist. The following lesions included in this study were scored on a scale of 0 to 4 from low to high severity: inflammation, hepatocellular degeneration, hepatocellular necrosis, foci of hepatocellular alteration, and Ito (stellate) cell hyperplasia.

### Quantification and Statistical Analyses

Statistical analysis was performed using GraphPad Prism software^20^. Tumor multiplicity, sequence rearrangement mutation frequencies, *gpt* mutant fractions, cleaved caspase-3 staining, and histopathology scores were compared by Mann-Whitney *U*-test. γH2AX staining and MN frequency were compared with unpaired Student’s *t*-test. A *P* value was considered significant if less than 0.05.

## RESULTS

We previously studied mutations, DNA damage, cell death, histopathological changes, and cancer in livers of WT, *Aag*-deficient (*Aag^-/-^*), and *Aag*-overexpressing (*AagTg*) mice following NDMA exposure^20^. Specifically, mice were exposed twice to NDMA, once on postnatal day 8 (3.5 mg/kg) and once on day 15 (7 mg/kg), and we monitored carcinogenic molecular, cellular, and physiological changes as they unfolded over time^20,37^. Here, we have expanded the results published in Kay et al., 2021 by specifically reporting on the impact of sex on NDMA-induced biological responses.

### Cancer

Previously, we showed that *Aag*^-/-^ mice were significantly more prone to NDMA-induced liver cancer 10 months post-exposure compared to WT, and that overexpression of *Aag* suppressed cancer^20^. Upon classifying these data by sex, we consistently found that males were more prone to cancer than females. For WT mice exposed to NDMA, 80% of males developed liver tumors, compared to just 46% cancer incidence in females (Table 1). Results also suggested an increase in the number of NDMA-induced tumors per liver for males compared to females (p = 0.05; Table 1 and Figure 1), which may reflect accelerated tumorigenesis.

**Table 1:** Tumors per mouse and cancer incidence.

| Genotype | Sex | Median # of tumors/mouse | Incidence (% mice with tumors) |
| --- | --- | --- | --- |
| WT | M | 1 | 12/15 (80%) |
|  | F | 0 | 6/13 (46%) |
| *Aag<sup>-/-</sup>* | M | 8 | 13/13 (100%) |
|  | F | 1 | 9/11 (81%) |
| *AagTg* | M | 1 | 8/12 (67%) |
|  | F | 0 | 2/12 (17%) |

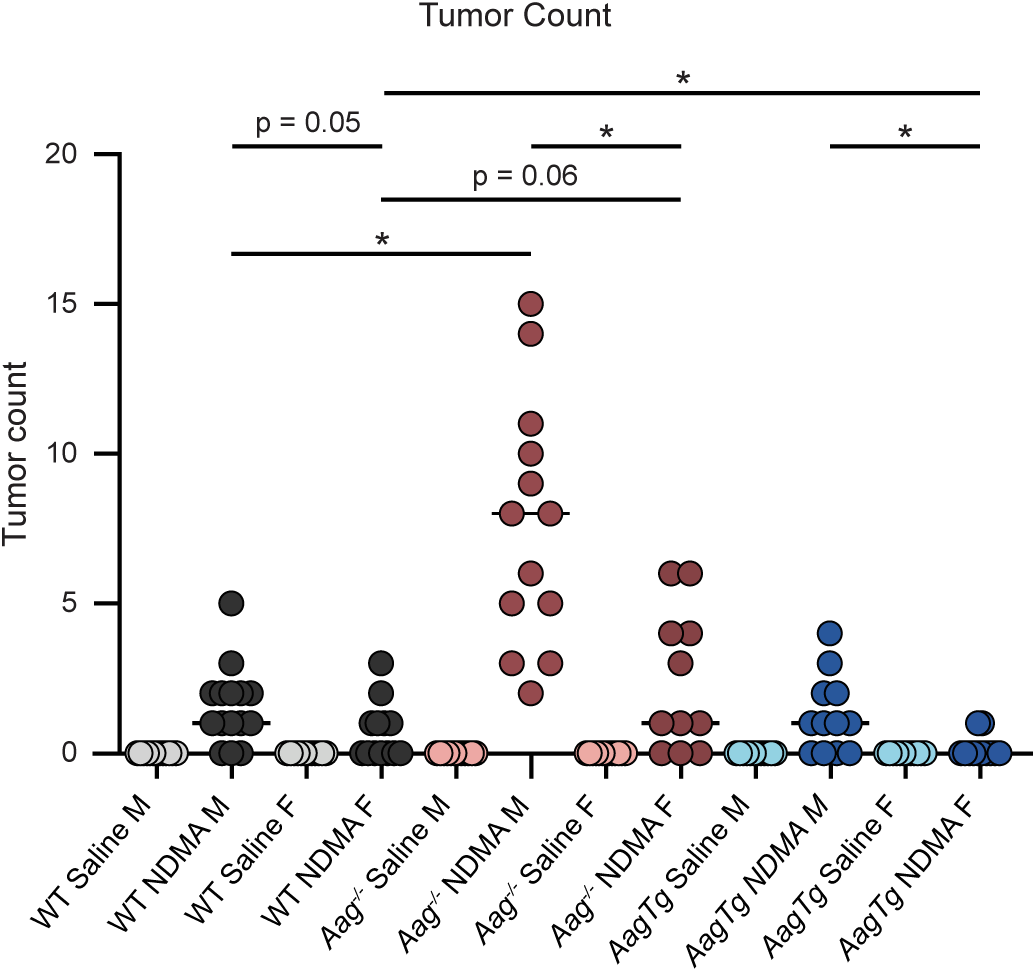

The sex-related difference in cancer susceptibility was more apparent in *Aag^-/-^* mice. The difference between males and females was particularly pronounced when considering the number of tumors. Males harbored significantly more NDMA-induced tumors than females, with a median of 8 liver tumors per male mouse compared to just 1 tumor per female (Table 1 and Figure 1). Similar to WT mice, cancer incidence also appeared to be reduced in females, though the results did not reach statistical significance (Table 1).

We also previously showed that overexpression of AAG suppressed NDMA-induced cancer incidence compared to WT^20^. Separating these data by sex, we found suppression of NDMA-induced tumors by AAG overexpression was evident in female mice. Only 17% of NDMA-exposed *AagTg* females developed macroscopic tumors, compared to 67% of *AagTg* males (Table 1). Tumor counts were also significantly lower in *AagTg* females (median = 0) versus males (median = 1; Table 1 and Figure 1). In fact, *AagTg* females were so resistant to liver cancer that the difference in tumor multiplicity between saline- and NDMA-treated *AagTg* females was not statistically significant, unlike every other saline-NDMA comparison.

We also found uniformity in the combination of sex and genetic factors associated with increased and decreased cancer susceptibility when we compared *Aag*^-/-^ and *AagTg* cohorts to WT mice. Consistent with the increased susceptibility to NDMA-induced liver tumors among males and among *Aag*^-/-^ mice, *Aag*^-/-^ males developed significantly more tumors compared to WT males, whereas the difference between tumor multiplicity in WT and *Aag*^-/-^ females did not reach statistical significance (p = 0.06) (Figure 1). Similarly, tumor multiplicity was significantly reduced in NDMA-treated *AagTg* females compared to WT females while this difference was not statistically significant for *AagTg* males versus WT males.

Altogether, these data demonstrate sex bias in NDMA-induced tumors, with male mice displaying increased vulnerability irrespective of genotype. Given that mutagenesis was strongly predictive of cancer induction, we next evaluated male-female differences in mutations.

### Mutations

Our experimental mice were homozygous for two transgenic reporter genes, *RaDR* and *gptD*, that allow for measurement of sequence rearrangement and point mutations in mouse livers, respectively. Briefly, RaDR mice contain two expression cassettes for enhanced green fluorescent protein (EGFP), each lacking critical sequence information either on the 5’ end or the 3’ end of the cDNA such that the protein products do not fluoresce^38^. HR with misalignment between the repeats can restore full length sequence, leading to the expression of fluorescent EGFP and enabling quantification of *de novo* recombination events *in situ* with fluorescence microscopy. The *gptD* transgene contains λ phage that when mutated, packaged, and infected into *E. coli* confers resistance to 6-thioguanine^39,40^. The frequency of 6-thioguanine resistant-colonies reflects the number of mutant transgenes present in the tissue. Sequence rearrangement and point mutations are important drivers of cancer, and the *RaDR^R/R^;gpt^g/g^*transgenic mouse model uniquely enables measurement of both mutation types in the same tissue from a single animal.

We have previously demonstrated that the RaDR substrate can detect spontaneous recombination events in mouse liver^38^, which we found to be elevated in *Aag^-/-^* mice compared to WT^20^. Interestingly, here we found that even in the absence of NDMA, rearrangement mutations in the saline-treated controls were significantly elevated in males compared to females, and this was the case across all three genotypes (Figure 2A; significance markers not shown in figure for ease of reading).

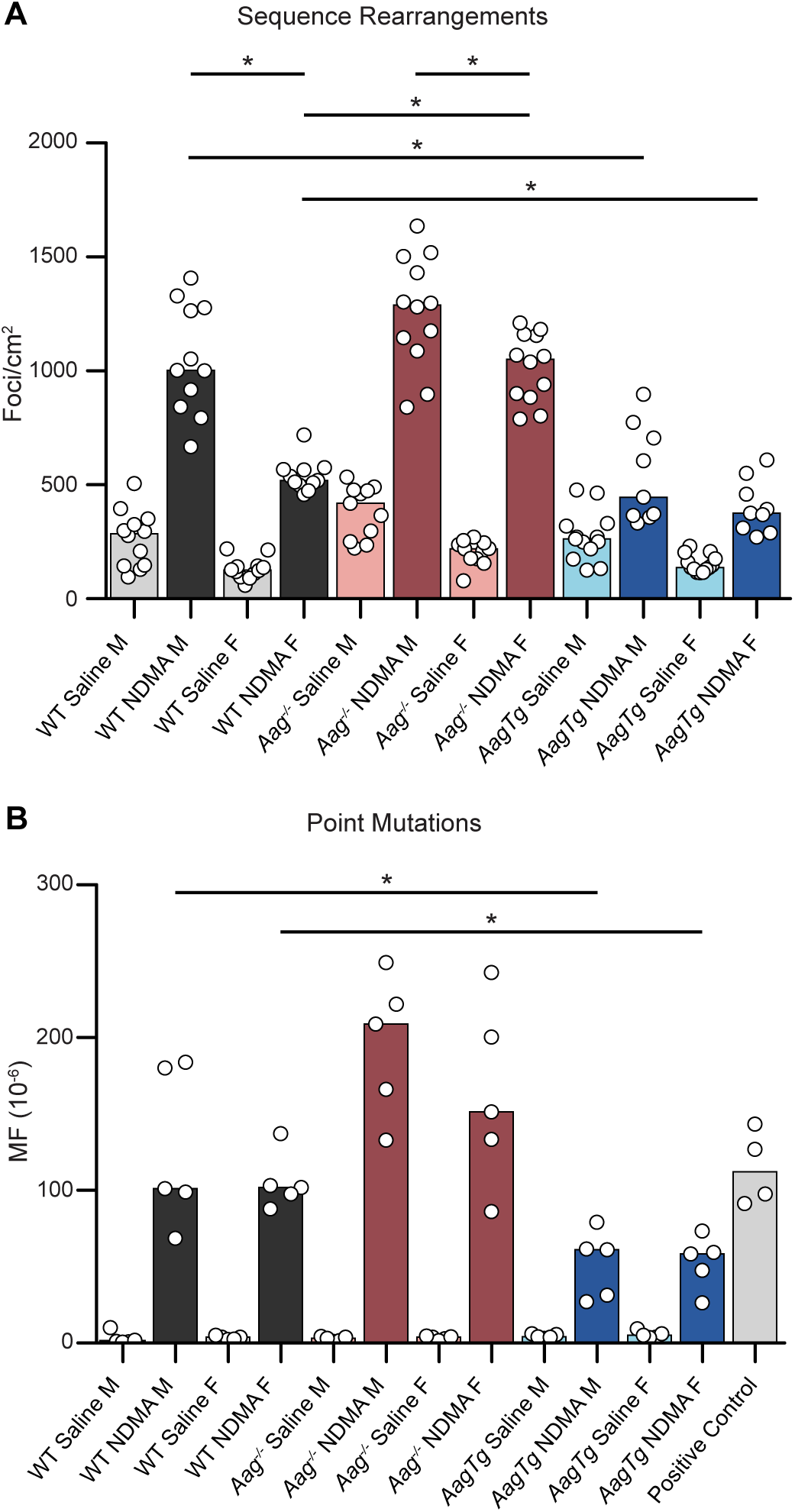

We had also shown that NDMA exposure significantly enhanced the levels of mutagenic recombination events in the liver 10 weeks after exposure and that AAG activity was protective against these mutations^20^. Separating the results by sex revealed a strong bias, with significantly higher frequencies of large-scale mutations in NDMA-treated males compared to females among WT and *Aag*^-/-^ mice, but not *AagTg* mice (Figure 2A). We also found that the loss of AAG increased susceptibility to NDMA-induced mutations to a greater extent in females (*Aag*^-/-^ females versus WT females p < 0.0001) than in males (*Aag*^-/-^ males versus WT males p = 0.06; Figure 2A). Consistent with our previous findings that *Aag* overexpression suppressed NDMA-induced sequence rearrangements, both *AagTg* males and females were significantly protected from NDMA-induced sequence rearrangements compared to WT (Figure 2A).

In addition to large-scale rearrangements, NDMA exposure in mice also generated point mutations^20,35^. We previously found that NDMA treatment induced significantly more point mutations in *Aag^-/-^* mice and significantly fewer point mutations in *AagTg* mice compared to WT^20^. Upon separating these data by sex, *AagTg* mice still showed fewer point mutations than WT, though the differences between *Aag^-/-^* and WT were not statistically significant (Figure 2B). Note that significant sex-related differences in NDMA-induced point mutations were not observed within any genotype, although there was a trend toward reduced susceptibility to point mutations in the female *Aag*^-/-^ mice compared to males (Figure 2B). A limitation to the point mutation analysis is the high mouse-to-mouse variation, which contributes to the lack of statistically significant differences.

### DNA Damage

In our prior study, we analyzed the genotoxicity of NDMA by γH2AX staining and MN quantification at 24 hours and 48 hours post-NDMA exposure, respectively^20^. Depending on the morphology of γH2AX staining, different biological consequences can be inferred. Specifically, punctate subnuclear γH2AX foci are indicative of DNA strand breaks, pan-nuclear γH2AX staining occurs in response to overwhelming DNA damage, and super-bright pan-nuclear γH2AX staining is thought to reflect toxic replication stress during S-phase^20^. MN can form during mitosis as a result of fragmented chromosomes.

Our earlier findings showed that NDMA led to an increase in the frequency of cells with ≥ 5 γH2AX subnuclear foci as well as pan-nuclear γH2AX staining^20^. The NDMA-induced increase in γH2AX foci relative to saline controls was largely preserved upon separating the data by sex (Figure 3A). In addition, *Aag^-/-^* mice tended to have lower induction of γH2AX foci and pan-nuclear staining relative to WT and *AagTg*^20^, which was only statistically significant for γH2AX foci in *Aag^-/-^* females compared to WT females when the data were stratified by sex (Figure 3A-B). There were no significant differences between males and females for either γH2AX foci or pan-nuclear staining within any genotype (Figure 3A-B).

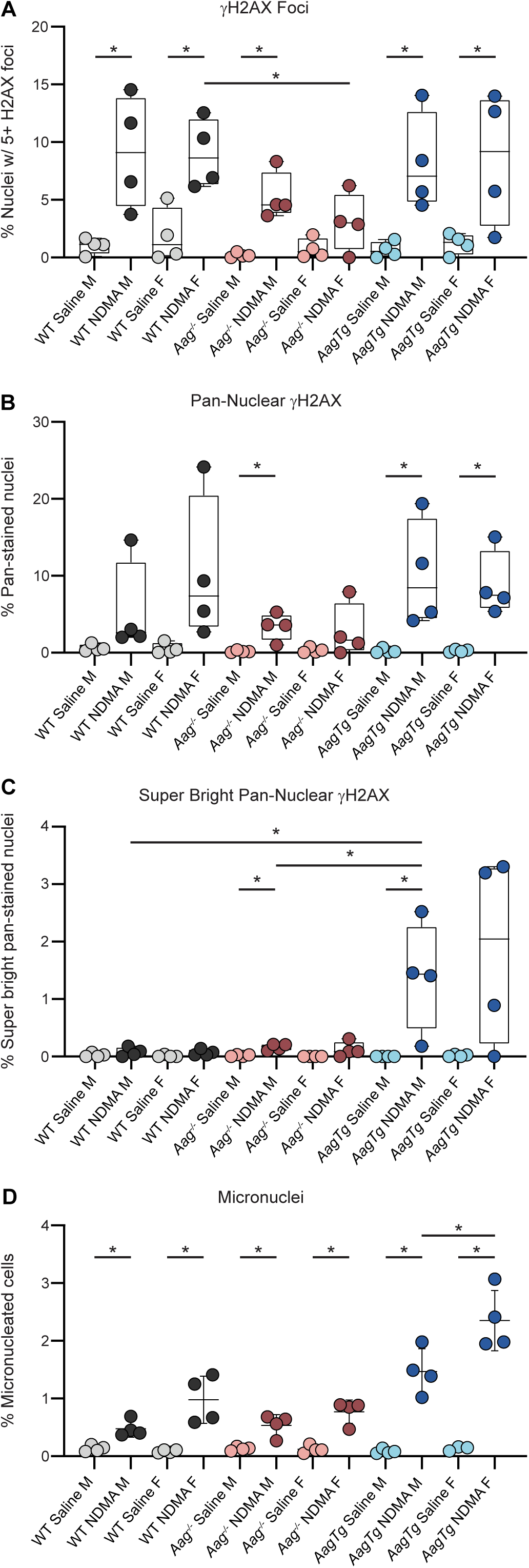

With regard to NDMA-induced super-bright pan-nuclear γH2AX staining, we previously observed a substantial increase specifically for *AagTg* mice relative to both WT and *Aag^-/-^* mice^20^. Consistent with this observation, after splitting the data for *AagTg* by sex, there was still an apparent increase in super-bright cells relative to saline, though not statistically significant for *AagTg* females. Compared to WT, *AagTg* males had more super-bright cells, and though the same trend is apparent in females, it did not reach statistical significance (p = 0.08), presumably due to high interindividual variation (Figure 3C). While the *Aag^-/-^* males showed a statistically significant increase in NDMA-induced super-bright pan-nuclear stained cells compared to saline, the magnitude of the effect was quite small (Figure 3C).

Consistent with sex-pooled data^20^, there was a significant induction of MN by NDMA upon separating the data by sex for all genotypes, with *AagTg* mice displaying the highest levels (Figure 3D). Sex-separated data also revealed an interesting trend of increased susceptibility to NDMA-induced MN in females compared to males, with a statistically significant difference in the *AagTg* mice (Figure 3D). Overall, the only significant difference observed when comparing the levels of DNA damage (either γH2AX or MN) between males and females was the increased susceptibility of *AagTg* females to NDMA-induced MN.

### Toxicity

We previously reported that *Aag*^-/-^ livers were more susceptible to apoptosis, necrosis, and inflammatory cell infiltration compared to WT, and that *AagTg* livers experienced even more severe cytotoxicity and inflammation^20^. We also observed lethality in NDMA-treated *AagTg* mice, wherein over 12% of *AagTg* mice died within two weeks of exposure^20^. Given that others have shown that males and females differ in susceptibility to inflammation, including in the liver^30,41,42^, we anticipated that males would show increased toxicity phenotypes over females. Instead, however, the levels of apoptotic cells and histopathological scores did not differ between males and females 24 hours post-NDMA exposure (Supplemental Figure 1), and the same proportion of *AagTg* males and females suffered acute lethality (Supplemental Table 1).

### Tissue Pathology

To gain additional information as to why male and female mice differ in tumor incidence, we analyzed livers from mice 10 months post-NDMA exposure for hepatocellular degeneration, which can be caused by a variety of toxicants including xenobiotics^43,44^. Indeed, NDMA significantly promoted degeneration in WT and *AagTg* mice compared to saline controls (Figure 4A). Remarkably, male mice appeared to be more susceptible to this type of liver injury for all genotypes (Figure 4A), though this comparison did not reach statistical significance for *Aag^-/-^*(p = 0.09). These findings are consistent with prior literature pointing to livers of male mice being more susceptible to liver damage^30^.

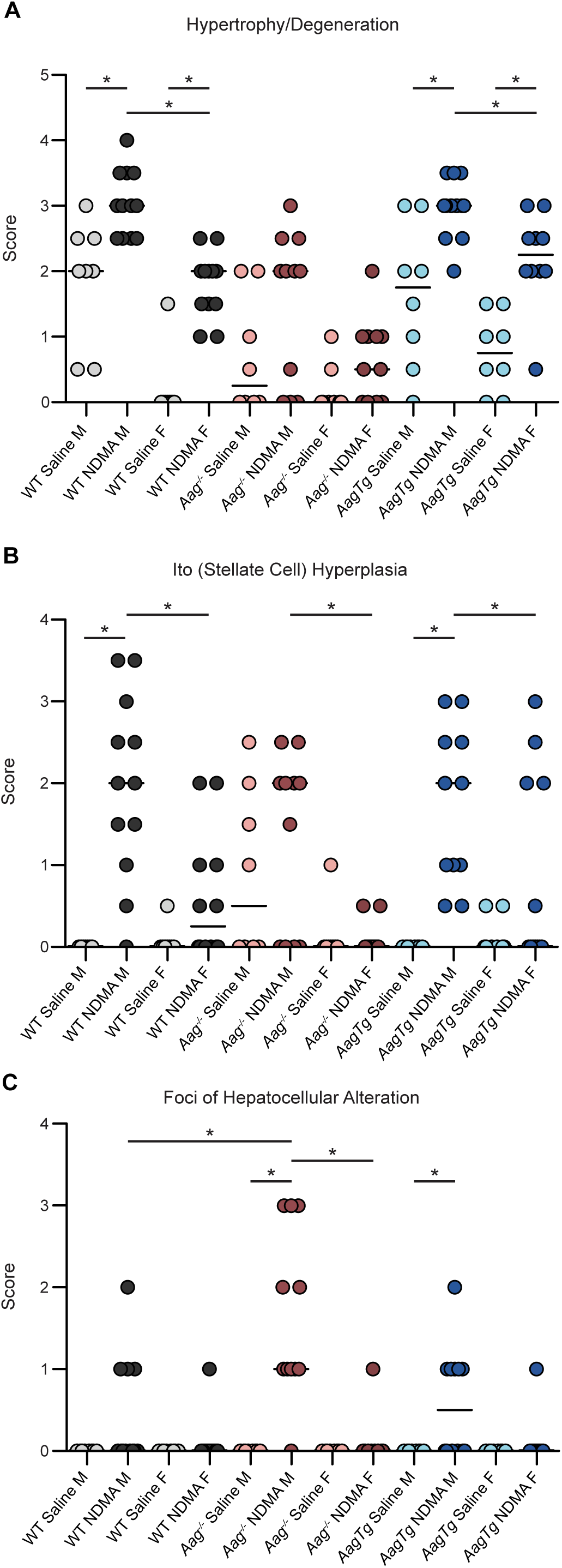

Histopathological analyses of livers from mice 10 months post NDMA exposure also showed that males were significantly more susceptible than females to Ito cell hyperplasia across all genotypes (Figure 4B). Interestingly, induction of Ito cell hyperplasia by NDMA was only statistically significant for WT males and *AagTg* males relative to saline controls. Ito hyperplasia refers to the proliferation of Ito cells (a.k.a., stellate cells). Ito cell proliferation can lead to increased fibrosis, ultimately contributing to liver disease^45^. In all, observations of hepatocellular degeneration and Ito cell hyperplasia clearly showed that males were more susceptible to liver damage than females.

Finally, we analyzed foci of hepatocellular alterations, which are aggregates of hepatocytes with changed morphology. There is a correlation between hepatocellular alterations and cancer, and it is thought that hepatocellular alterations precede tumor formation^46^. *Aag^-/-^*male mice displayed the greatest increase in NDMA-induced hepatocellular alterations, which was significantly elevated in comparison to WT males (Figure 4C). Consistent with the increase in susceptibility to liver tumor formation (Table 1 & Figure 1), males were generally more susceptible to NDMA-induced hepatocellular alterations compared to females, though this was only statistically significant in *Aag^-/-^*mice (Figure 4C). Altogether, tissue pathology at 10 months post-NDMA exposure revealed enhanced liver damage in male mice compared to females, which may be an important mechanism underlying the observed sex bias in NDMA-induced liver cancer.

### Integrated Analysis of Responses to NDMA Exposure

We integrated phenotypic observations over time using radar plots (Figure 5). For each endpoint, results are normalized to the highest scoring group. For example, NDMA-treated *Aag*^-/-^ males had the highest median number of tumors, so all tumor burden medians were normalized to NDMA *Aag*^-/-^ males; similarly, *AagTg* NDMA females had the highest mean induction of MN, so all MN means were normalized to *AagTg* NDMA females. Although differences in some endpoints were not statistically significant, the radar plots provide the ability to see overall trends holistically. As we found with sex-pooled data^20^, *Aag*^-/-^ mice were more susceptible to long-term adverse effects (mutations and cancer, on the left) whereas *AagTg* mice experienced higher initial toxicity (DNA damage and lethality, on the right) but fewer persistent effects. Importantly, males were more vulnerable to persistent mutations and eventual cancer, while females were somewhat more sensitive to early toxicity, although the increased sensitivity for females was only statistically significant for MN induction.

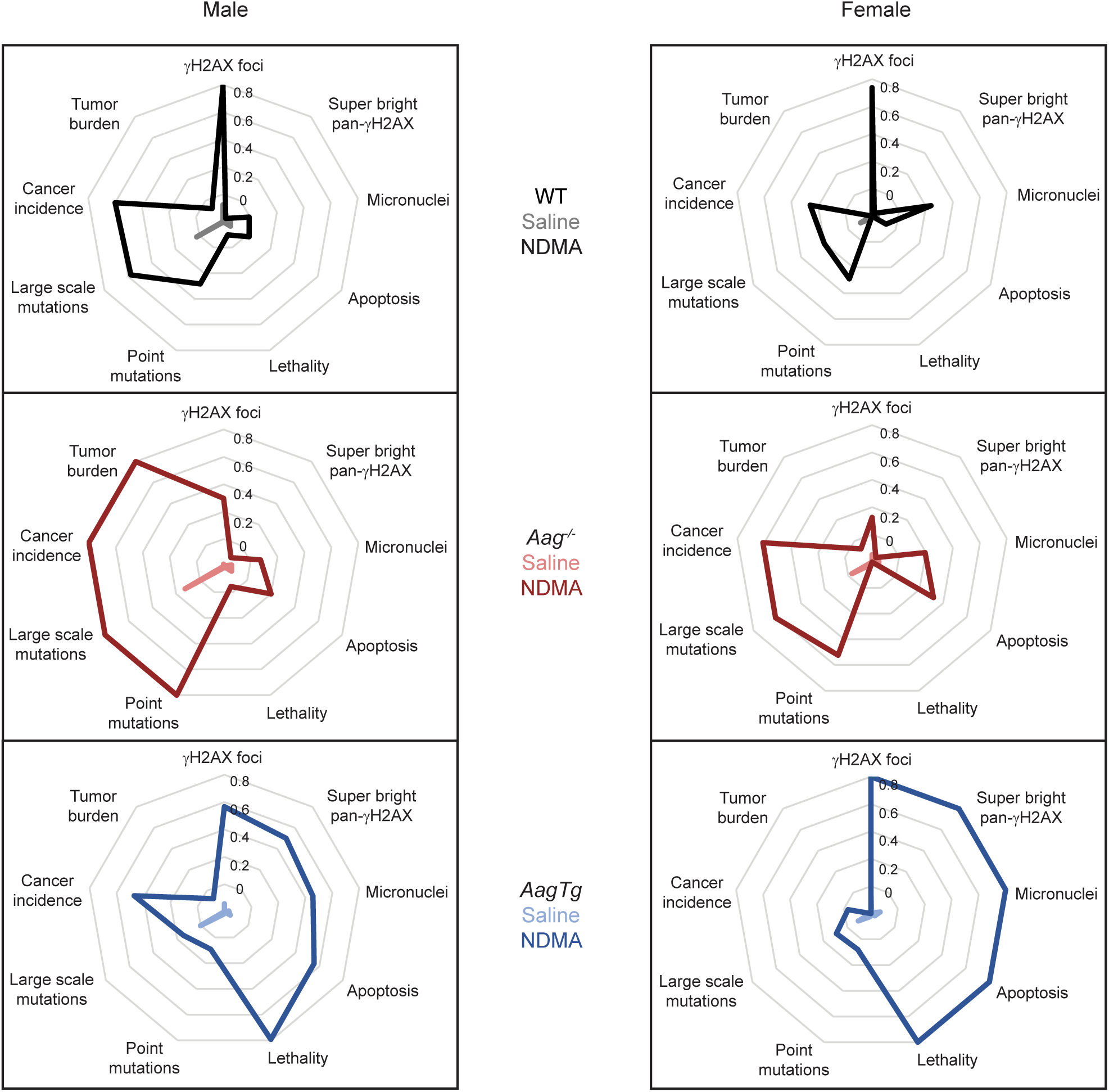

## DISCUSSION

It is well established that males are more susceptible to liver cancer than females, which has been observed in both humans and mice^18,29,30,32–34,47–49^. Furthermore, there are sex-related differences in the effect of environmental agents as triggers of this disease^31^. Indeed, prior research focusing on the impact of nitrosamines on liver cancer in mice has shown that males are more vulnerable than females^18,32–34^. Here, we have expanded upon that prior work to delve into the sex-specific molecular, cellular, genetic and physiological changes that occur with NDMA exposure, which provides insights into the mechanisms underlying sex bias in NDMA-induced cancer. Overall, we found that males were not only more prone to NDMA-induced cancer (Table 1 & Figure 1), but they were also more prone to sequence rearrangements ten weeks post-exposure (Figure 2A) and liver pathology ten months post-exposure (Figure 4). In contrast, signs of early damage were slightly higher in the females (Figure 5).

Across all genotypes, males were more susceptible to NDMA-induced liver cancer than females (Table 1 and Figure 5). The most striking sex-related enhancement in tumor incidence was in *AagTg* mice, while *Aag*^-/-^ mice showed the greatest sex-related enhancement in tumor multiplicity (Table 1 and Figure 1). By combining the variables of sex and genotype, it becomes clear that maleness combined with *Aag* deletion creates the most conducive conditions for tumor development, whereas femaleness combined with *Aag* overexpression is most protective against cancer.

We saw a strong association between the relative susceptibility to sequence rearrangements (Figure 2A) and the relative susceptibility to cancer (Figure 1 and Table 1). This observation was consistent with the pooled data from males and females wherein sequence rearrangements, which occur relatively soon after exposure, are highly correlated with downstream cancer susceptibility^20^. Approximately 50% of the genome is composed of repetitive sequences^50,51^, so a propensity for recombination events at the HR substrate in the RaDR locus likely reflects a biologically relevant mechanism for carcinogenic large-scale rearrangement mutations.

An important contribution of this study is the identification of male sequence rearrangement mutation bias in somatic tissue. Until now, male mutation bias has primarily been shown in germline^52–54^ or cancer contexts^47–49,55^. However, these earlier studies often do not specify the type or mechanism of mutation (*e.g*., point mutations, frameshifts, sequence rearrangements), and they have not analyzed physiologically normal somatic tissues. A remarkable finding in comparing sex differences in mutagenesis in the present work was that, for all genotypes, females accumulated significantly fewer spontaneous sequence rearrangement mutations compared to their male counterparts (Figure 2A). Additionally, among NDMA-treated mice, WT and *Aag^-/-^* females developed fewer sequence rearrangements than males. To our knowledge, this is the first demonstration that males are more prone to HR-driven sequence rearrangement mutations in the liver both spontaneously and following exposure to a carcinogenic alkylating agent, pointing to a novel mechanism underlying the known increase in susceptibility to liver cancer for males.

The fact that HR-driven sequence rearrangements are lower in females than males (Figure 2A) may be related to the fact that estrogen receptor α (ERα) facilitates the two major pathways that repair NDMA-induced DNA damage: BER initiated by AAG (which removes the replication-blocking 3MeA lesion) and direct reversal by *O*^6^-methylguanine-DNA-methyltransferase (MGMT, which removes the aberrant methyl group from *O*^6^MeG). ERα interacts directly with AAG to enhance excision activity^56^, thus reducing the potential for replication blockage by 3MeA to lead to mutagenic recombination events. Furthermore, the *Mgmt* promoter contains ERα responsive elements resulting in enhanced MGMT expression following estrogen exposure^57^. In addition, spent MGMT negatively regulates ERα-mediated cell proliferation, preventing cell division before repair of *O*^6^MeG is complete^58^. While the *O*^6^MeG lesion does not block replication, mismatch repair futile cycling leads to persistent single strand breaks, which can be recombinogenic^59,60^. Thus, even in *Aag*^-/-^ mice, which do not have estrogen-enhanced excision of 3MeA, estrogen is likely protective against mutagenic recombination through enhanced MGMT expression (reducing futile cycling) and negative regulation of cell division (by spent MGMT).

Interestingly, previous studies in mammary cell lines have shown that estrogen delays γH2AX resolution, reduces HR, and induces chromosomal instability following irradiation^61^, which is consistent with our observations of increased MN (Figure 3D) and fewer sequence rearrangements (Figure 2A) in females. The greater induction of MN in WT and *AagTg* females may be associated with HR inhibition by estrogen, preventing resolution of strand breaks and leading to chromosomal fragments during mitosis. Another possible explanation for increased MN induction in WT and *AagTg* females is that the interactions between ERα and AAG may enhance glycosylase excision^56^, thus leading to increased BER-intermediate strand breaks that result in double-strand breaks during replication. We hypothesized in our previous publication that the excess BER-intermediate strand breaks in NDMA-exposed *AagTg* mice reduced mutations through cytotoxicity^20^. However, while NDMA-treated females displayed higher induction of MN (Figure 3D), they did not show overt differences in toxicity (Supplemental Figure 1 & Supplemental Table 1). Thus, increased clearance of damaged cells likely does not explain the reduced recombinant cells in females. Interestingly, results from the Samson laboratory previously showed that in the kidney, retina, and cerebellum, cytotoxicity of alkylating exposures is reduced in females, which suggests tissue-specific differences in the impact of *Aag* on cytotoxicity^62–64^.

Long-term morphological changes and liver damage were more apparent in NDMA-exposed males versus females (Figure 4). In nearly all cases, the levels of NDMA-induced hepatocellular degeneration, Ito hyperplasia, and foci of hepatocellular alteration were elevated in males. The liver is a sexually dimorphic organ, and hormones play an integral role in modulating responses to environmental insult. For example, one study showed that exposure to diethylnitrosamine in mice significantly increased serum interleukin-6 (IL-6) levels in males compared to females, which was suppressed by treatment with an estrogen receptor agonist^32^. IL-6 is an inflammatory cytokine primarily produced by Kupffer cells in the liver, and it can promote survival, proliferation, invasion, angiogenesis, and metastasis through the IL-6/STAT3 axis^30^. Interestingly, knockout of IL-6 abolished the sex-related difference in diethylnitrosamine-induced liver tumor development^32^. The suppression of IL-6 in females is one of the many estrogen-mediated effects that has been shown to reduce the occurrence and aggression of HCC. In contrast, androgens in males have been found to promote proliferation and survival, which can lead to the retention of damaged cells and carcinogenesis^30^.

Our observations of differences in WT, *Aag*^-/-^, and *AagTg* males and females in response to NDMA reveal surprising patterns that warrant further investigation. In particular, we found remarkable protection against mutations and cancer in females, but no sex-related difference in γH2AX and an increase in NDMA-induced MN among *AagTg* females compared to males (Figure 3). Similar levels of apoptosis and necrosis among males and females (Supplemental Figure 1A and B) suggest the differences in mutations are not due to early clearance of damaged cells. It would be interesting to analyze a larger number of mice for γH2AX, MN, cytotoxicity, and point mutations to determine if slight sex-related differences exist that were not apparent here. Additionally, since estrogen is produced at similar levels in neonatal males and females^65^, it may also be informative to analyze DNA and tissue damage at a later timepoint when sex hormone production is significantly different. This could help to elucidate possible windows of susceptibility to NDMA exposure for both sexes, a topic of great importance for toxicological risk analysis.

Taken together, we demonstrate here the importance of both AAG-mediated DNA repair and sex in response to NDMA exposure. Key findings from this study reveal that male mice are more susceptible to NDMA-induced large-scale mutations and long-term liver damage than females, helping to explain the increased susceptibility to liver cancer observed among men.

## ACKNOWLEDGEMENTS

This work was supported by the National Institutes of Health (NIH)/National Institute of Environmental Health Sciences (NIEHS) Superfund Basic Research Program Grant P42-ES027707, NIEHS Core Center Grant P30-ES002109, NIEHS Toxicology Training Grant T32-ES007020, NIH Grant R01-CA080024, NIEHS grant K01-ES036182, NIEHS Small Business Innovation Research Grant R44-ES0264644, NIH Biomedical Technology Research Resource Grant 5-P41EB015871-32, and the Center for Advanced Imaging at Harvard University. We thank Caroline Atkinson and Joanna Richards in the MIT Division of Comparative Medicine for H&E and cleaved caspase-3 staining and Kathleen Cormier in the MIT Koch Institute Histology Core for Aperio slide scanning of caspase-stained slides. We appreciate the assistance of Aimee Moise in necropsies and RaDR imaging and Judy Yau in gpt analysis. This content is solely the responsibility of the authors and does not necessarily represent the official views of the NIH.

---

**Supplemental Table 1:**

| Genotype | Sex | Total animals injected with NDMA | Total dead within two weeks | Premature deaths (% injected) |
| --- | --- | --- | --- | --- |
| WT | M | 75 | 1 | 1.3 |
|  | F | 62 | 0 | 0 |
| *Aag<sup>-/-</sup>* | M | 50 | 1 | 2 |
|  | F | 43 | 0 | 0 |
| *AagTg* | M | 55 | 7 | 12.7 |
|  | F | 47 | 6 | 12.8 |

