## Supplementary figures and images for "Sex and Alkyladenine DNA Glycosylase Expression are Key Susceptibility Factors for NDMA-induced Mutations, Toxicity, and Cancer"

### Supplemental Figure 1

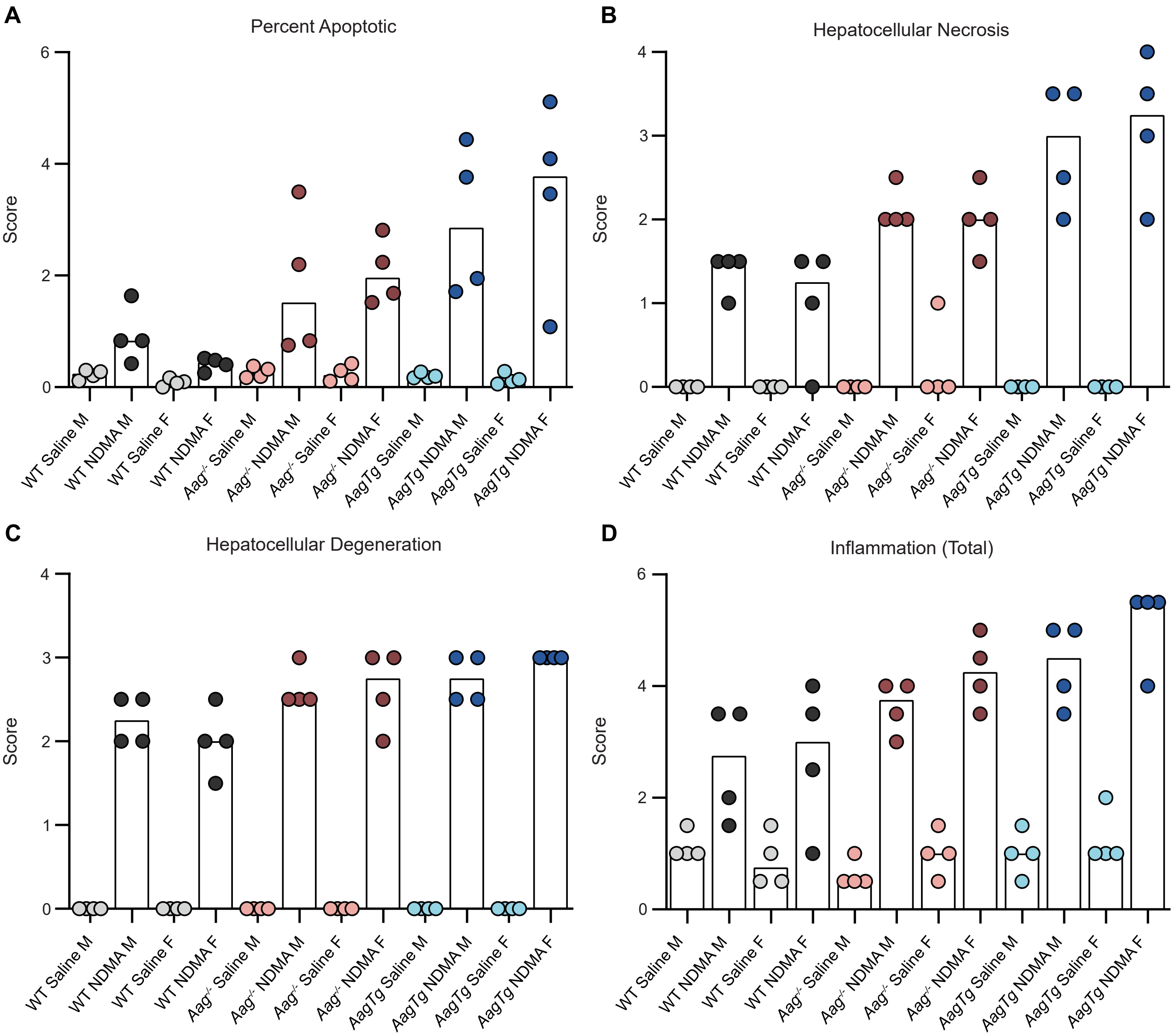
